# The anticipation and perception of affective touch in women with and recovered from Anorexia Nervosa

**DOI:** 10.1101/2020.02.23.961367

**Authors:** Laura Crucianelli, Benedetta Demartini, Diana Goeta, Veronica Nisticò, Alkistis Saramandi, Sara Bertelli, Patrizia Todisco, Orsola Gambini, Aikaterini Fotopoulou

## Abstract

Disruptions in reward processing and anhedonia have long being considered as possible contributors to the aetiology and maintenance of Anorexia nervosa (AN). Recently, interoceptive deficits have also been observed in AN, including reduced tactile pleasure. However, the extent to which this tactile anhedonia is specifically liked to an impairment in a specialized, interoceptive C tactile system originating at the periphery, or a more top-down mechanism in the processing of pleasant tactile stimuli remains debated. Here, we investigated two related hypotheses. First, we examined whether the differences, between patients with AN and healthy controls in the perception of pleasantness of touch stimuli delivered in a CT-optimal manner versus a CT non-optimal manner would also be observed in patients recovered from AN. This is important as tactile anhedonia in acute patients may be the secondary result of prolonged malnutrition, rather than a deficit that contributed to the development of the disorder. Second, we examined whether these three groups would also differ in their top-down, anticipatory beliefs about the perceived pleasantness of different materials touching the skin, and to what degree such top-down beliefs and related impairments in alexithymia and interoceptive sensibility would explain any differences in perceived tactile plesantness. To this end, we measured the anticipated pleasantness of various materials touching the skin and the perceived pleasantness of light, dynamic stroking touches applied to the forearm of 27 women with AN, 24 women who have recovered and 30 healthy controls using C Tactile (CT) afferents-optimal (slow) and non-optimal (fast) velocities. Our results showed that both clinical groups anticipated tactile experiences and rated delivered tactile stimuli as less pleasant than healthy controls, but the latter difference was not related to the CT optimality of the stimulation. Instead, differences in how CT optimal touch were perceived were predicted by differences in top-down beliefs, alexithymia and interoceptive sensibility. Thus, this study concludes that tactile anhedonia in AN is not the secondary result of malnutrition but persists as a trait even after otherwise successful recovery of AN and also it not linked to a bottom-up interoceptive deficit in the CT system, but rather to a learned, defective top-down anticipation of pleasant tactile experiences.

## 1. Introduction

Anorexia Nervosa (AN) is an eating disorder characterized by restriction of energy intake, by either avoiding food consumption (particularly in the “restricting-type” of AN), or purging (particularly in the “binge-eating/purging-type”), intense fear of gaining weight, disturbances in the experience of one’s own body weight or shape and unawareness of such disturbances (DSM-5, 2013). The associated malnutrition can have both cognitive (e.g. altered judgment and decision making) and physiological consequences (e.g. amenorrhea, reviewed by Zakzanis, Campbell & Polsinelli, 2010). Although several psychological and pharmacological treatments have been developed in the past two decades, outcomes in AN remain poor, with high rates of attrition, and frequent relapse (Carter, Blackmore, Sutandar-Pinnock, & Woodside, 2004; Steinhausen, 2002). Despite advances in the understanding of the heterogeneity of AN, it is still unclear whether some dysfunctions, such as serotonin (Haleem, 2012; Kaye, Fudge, & Paulus, 2009), or dopamine abnormalities (Avena & Bocarsly, 2012; Kaye, Wierenga, Bailer, Simmons, & Bischoff-Grethe, 2013; Scheurink, Boersma, Nergardh, & Sodersten, 2010) are a cause or consequence of malnutrition. It is also unknown whether individuals with anorexia nervosa have a primary disturbance of appetite regulation (including sensory, hedonic and motivational aspects of feeding) or whether pathological feeding behavior is secondary to other phenomena, such as an obsessional preoccupation with body image. For example, there are different and conflicting theoretical hypothesis about the precise role of dopamine in AN, with some suggesting that food restriction itself is rewarding in AN, while others indicate that the more general lack of sensory pleasure experienced by patients with AN contributes to their ability to forgo the pleasure of food (Keating, 2010; Kontis & Theochari, 2012; Walsh, 2013). Given this uncertainty regarding the rewarding value of food in AN, several studies have used monetary reward as a way to understand the mechanisms underlying patients’ response to and learning from reward and punishment (e.g. Decker et al., 2015; Bernardoni et al., 2018). However, it has become apparent that performance in such generic tasks can be differentially influenced by various physiological parameters (e.g. Fisher & Rangel, 2014; Wierenga et al., 2015) that need to be taken into account in AN studies.

Indeed, abnormalities have been found in several physiological domains, including altered subjective and neural responses to physiological sensations, such as taste (Oberndorfer et al., 2013; Wagner et al., 2008; Cowdrey et al., 2011), hunger signalling (Monteleone, P. & Maj, 2013), heartbeat detection (Pollatos et al., 2008; Kerr et al., 2016), aversive breathing load (Berner et al., 2017), pain (Strigo et al., 2015) and affective touch (Crucianelli et al., 2016; Davidovic et al., 2018). These are domains of interoception, defined as the ability to sense and process the physiological condition of one’s body (Craig, 2002). Interoception lies at the heart of the physiological components of emotion (Craig, 2009) and forms the core of the embodied basis of self-representation (Garfinkel et al., 2013). Accordingly, it has been proposed that in AN (Khalsa et al., 2015), as in several other psychopathologies (Paulus & Stein, 2006; 2010; Khalsa et al., 2015; 2018), there is a dysregulated ability to adequately predict, sense and modulate what is happening in the physiological state of the body, resulting in symptoms such as alexithymia (Shah et al., 2016), increased self-related worry and poor self-efficacy (Stephan et al., 2016), and dysfunctional learning about bodily states over time (van den Bergh, 2017).

There is however some evidence suggesting that differential modalities of interoception may have a distinct effect on body awareness (Crucianelli et al., 2018). Moreover, while some afferent signals are hedonically neutral (e.g. heart rate), others such as noxious or gentle skin stimulation tend to also convey hedonic and motivational signals. Of interest here, ‘affective touch’, hereafter referring to a specialised tactile modality of gentle, stroking touch, thought to be (at least partly, see Marshall et al., 2019) mediated peripherally by unmyelinated, slow-conducting C-tactile (CT) afferents (Löken et al., 2009) and to be centrally processed by an interoceptive pathway converging at the insular cortex (Morrison, 2016; Kirsch et al., 2020), has been associated with subjective sensations of skin pleasure (Löken et al., 2009) and related behavioural measures (Pawling et al., 2017).

Moreover, while in a first consideration gentle, affective pleasant touch may not appear as symptom-specific as some of the other tested modalities of interoception (e.g., gut monitoring, hunger), recent findings in the field suggests that CT-optimal affective may be uniquely relevant to one’s bodily self-representation (Gentsch et al., 2016; Ciaunica & Fotopoulou, 2017). Specifically, CT-optimal touch has been found to increase embodiment during multisensory integration tasks such as the rubber hand illusion (Crucianelli et al., 2013, 2018; Lloyd et al., 2013; van Stralen et al., 2014) and the enfacement illusion (Panagiotopoulou et al., 2017). Moreover, CT-optimal, affective touch has been found to enhance feelings of body ownership in right-hemisphere patients with delusions of disownership (Jenkinson et al., 2019).

These considerations have led to three independent investigations of the anticipation and perception of CT-optimal, affective touch in AN. First, our group showed that 25 patients with acute, restrictive-type AN perceived gentle skin stroking touch at CT-optimal velocities as less pleasant relative to 30 healthy controls (HC), while there was no such difference for CT-suboptimal velocities (Crucianelli et al., 2016), suggesting that individuals with AN have abnormalities in the perception of the pleasantness of CT-mediated, affective touch. Davidovic and colleagues (2018) provided further behavioural support for this conclusion, showing that 25 patients with AN rated CT-optimal touch as less pleasant than a group of matched healthy controls. Functional neuroimaging results were mixed. While they found no differences amongst the two groups in the activation of insular cortex when examining the contrast between the condition of gentle touch and no stimulation, they did observe a relative hypoactivation of the left caudate nucleus, part of the reward-relevant circuitry of the striatum, in individuals with AN.

A third study (Bischoff-Grethe et al., 2018) focused on 18 individuals recovered from AN (RAN; both types) with a functional neuroimaging (fMRI) paradigm that included an anticipation phase (cues of the location of impending tactile stimulus) in addition to behavioural ratings of affective touch pleasantness in two locations, forearm (containing CT fibers) and palm (not containing CT fibers). While some differences in how RAN individuals anticipate both CT and non-CT touch were observed in this study, contrary to the above studies, the RAN group did not rate the pleasantness of CT-touch as less pleasant than controls and there were not any CT-specific neuroimaging findings at the group level. When individual traits were taken into account, individuals with RAN and with higher harm avoidance scores had lower BOLD responses during forearm touch anticipation in right ventral mid-insula, while other clinical variables, such as lower historical BMI and longer illness duration led to altered insula BOLD responses during CT touch, but these clinical severity-related clusters did not overlap with any task related clusters. While these findings suggest that deficits in both the anticipation and the perception of CT optimal touch may relate to some of the features of AN, caution is required when individual differences in such a small group are tested. In conclusion, this study tested individuals RAN and did not confirm the findings of the previous two studies in the sense that RAN individuals did not differed from controls in their subjective and neural responses to CT-mediated touch.

The above discrepancies between the studies may relate to the different paradigms and samples used. In this study, we aim to address such differences and put forward a related, new hypothesis about the perception of CT-mediated, affective touch in AN. Firstly, we used a paradigm that assessed both the perception of CT-optimal touch and top-down beliefs about pleasantness. Specifically, we asked participants to think about being touched by soft or rough material on the skin and to provide estimates of pleasantness of such experience. Next, participants were asked to rate the pleasantness of different brush strokes delivered at different CT-optimal and CT-suboptimal velocities on their forearm while blindfolded. We expected patients with AN to anticipate and perceive tactile stimuli as less pleasant than controls and we expected their reduced pleasantness ratings to be explained by less differentiation in the pleasantness perception between CT-optimal and suboptimal tactile stimuli. We then investigated the relationship between the performance at the two tasks to explore to what extent anticipatory beliefs about tactile pleasantness, and related deficits in alexithymia and interoceptive sensibility in the AN group might influence the pleasantness of actual tactile experiences. Secondly, we recruited both acute and recovered AN individuals (restrictive type) and examined whether the above hypotheses applied also to recovered patients in comparison to healthy controls. Testing recovered, as well as acute patients in eating disorders research is of high importance as the malnutrition in the acute stages of illness can itself cause cognitive and emotional deficits and hence it is not possible to determine whether abnormalities in perceiving pleasure and other symptoms are a cause or a consequence of starvation. In brief, this study aimed to investigate (1) whether the anticipation and perception of pleasant touch would be reduced in patients who have recovered from AN as well as in patients with acute AN compared to healthy controls; (2) whether this tactile anhedonia will be best explained by deficits in the CT-afferent system, top-down beliefs about the pleasantness of affective touch, or their combination.

## 2. Methods

### 2.1. Participants

Twenty-seven female participants with restrictive type AN and twenty-four patients who have recovered from restrictive type AN (RAN) were recruited from the Day care Unit of St Paolo’s Hospital in Milan and from Comunità Villa Miralago in Cuasso al Monte (VA), and Casa di Cura Villa Margherita in Arcugnano, Italy. All patients were formally diagnosed (Diagnostic and Statistical Manual of Mental Disorders: Fifth Edition, DSM-V, American Psychiatric Association, 2013) by a psychiatrist. Inclusion criteria for the AN group, beyond AN restrictive type diagnosis, included a BMI below 18.5 (Table 1). As in previous studies, AN recovery was defined as being weight-restored (BMI above 18.5), regular menstrual cycles and no binge eating, purging, or restrictive eating patterns for at least 1 year before the study (e.g. Wierenga et al. 2015). Twenty-seven matched heathy controls (HC) of Italian nationality were also recruited. Exclusion criteria for the HCs included any history of psychiatric or neurological condition. Exclusion criteria for all the groups included being left-handed, being below 18 or above 45 years, having any medical conditions that result in skin conditions (e.g. psoriasis, eczema, etc) or any visible scar on the left forearm, and any substance abuse. Institutional ethics approval was obtained, and all participants provided written informed consent to participate. All participants received a €10 voucher, £10 in cash or University credits.

**Table 1.** Clinical profile of the Anorexia Nervosa (AN) and Recovered Anorexia Nervosa (RAN) groups.

|  | **AN** | **RAN** |
| --- | --- | --- |
| Years of Illness | 10.92 [10.23] | 4.18 [2.92] |
| Age of Onset | 17.23 [4.45] | 16.78 [2.7] |
| Years from Recovery | N/A | 2.97 [3.11] |
| Psychiatric comorbidities | 5 depression<br>2 bipolar disorder<br>8 anxiety disorder<br>2 OCD<br>4 Borderline PD<br>7 multiple comorbidities | 8 depression<br>8 anxiety disorder<br>3 OCD<br>2 Borderline PD |
| Psychiatric Treatment | 14 SSRIs<br>2 tranquilizers | 6 SSRIs<br>3 tranquilizers |
| Other comorbidities and treatment | 2 hypothyroidism<br>2 osteoporosis<br>dietary supplements |  |

### 2.2. Psychometric Assessments

All participants completed the subscale “interoceptive awareness” from the Eating Disorder Inventory-2 [EDI-2; (Garner, 1984; Thiel & Paul, 2006)]. This subscale focused on the subjective identification of bodily signals, such as hunger and satiety. Questions are rated on a six-point scale, ranging from 1 (never) to 6 (always). High scores indicate potential issue with interoceptive awareness. They also completed the Body Awareness Questionnaire (BAQ) and the Toronto Alexithymia Scale (TAS-20).

The BAQ is a self-report measure of interoceptive sensibility (Shield, Mallory & Simons, 1989), consisting in 18 items assessing the subject’s sensitivity to physiological body processes to be replied to on a 7-point likert scale (from 1 = “Not at all true of me” to “7 = Very true of me”). Four subscales were calculated: (i) Note Response or Changes in Body Process; (ii) Predict Body Reaction; (iii) Sleep-Wake Cycle; (iv) Onset of Illness.

The TAS-20 is a well-validated self-report measure of alexithymia (Bagby, Taylor, & Parker, 1994), where participants are asked to rate their agreement to 20 statements on a 5-point likert scale (ranging from 1 = “strongly disagree” to “5 = strongly agree”). A total score above 61 (out of a maximum of 100) is considered a cut-off score of alexithymia. Three sub-scores were calculated, assessing: (i) Difficult Identifying Feelings (DIF); (ii) Difficulties Describing Feelings (DDF); (iii) Externally-Oriented Thinking (EOT).

### 2.3. Experimental procedure

Upon arrival, participants were given the information sheet and they were asked to sign the consent form, before completing the questionnaires: demographic, EDI, BAQ and TAS. Next, participants were familiarised with the rating scale, ranging from 0 (not at all pleasant) to 100 (extremely pleasant). Participants were invited to use the entire scale and to say any number between 0 and 100. As part of the familiarisation phase, participants were asked to think about being touched by two soft (i.e. cotton and velvet) and two rough objects or material (i.e. thorn and sandpaper) on their forearm and to rate the hypothetical related pleasantness (i.e. “How pleasant would it be to be touched by ‘cotton’ on your forearm?”). The order of the materials was counterbalanced between participants. This allowed us to collect a measure of anhedonia prior to the completion of the affective touch task, and also to make sure that participants fully understood how to use the rating scale (Figure 1).

**Figure 1.**
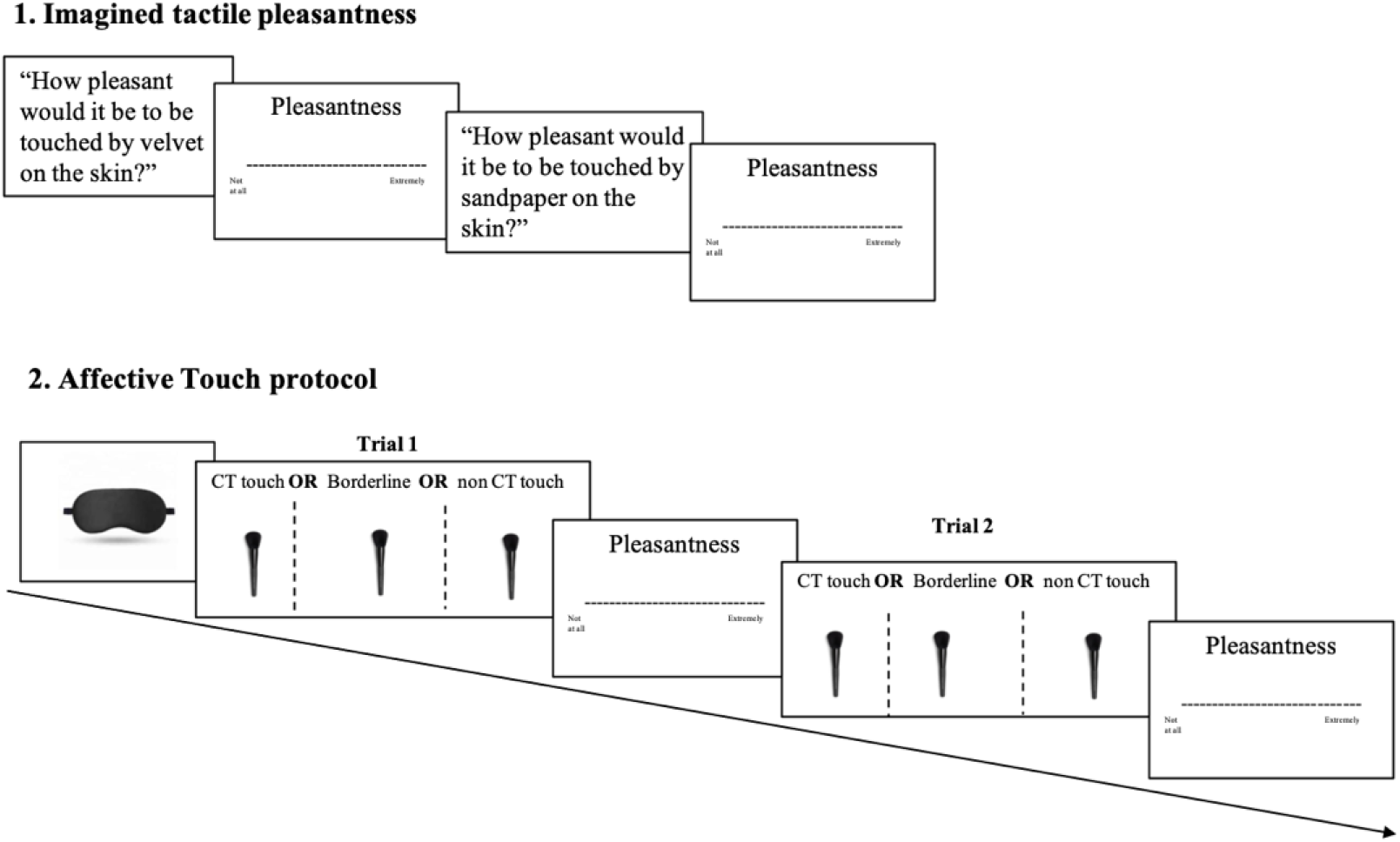
Schematic representation of the study procedure. All participants completed the same experimental procedure. 1. First, they were asked to answer four hypothetical questions about imagined touch, such as “How pleasant would it be to be touched by velvet on your skin?” using the VAS scale ranging from 0, not at all, to 100, extremely pleasant. 2. Participants were then asked to put on a blindfold before the experimenter delivered the touch on the left forearm (dorsal part) at CT-optimal, borderline or non-CT optimal velocities (in randomized order). Each touch lasted 3 seconds and was repeated 3 times for each velocity for a total of 15 trials. After each touch, participants were asked to rate the pleasantness of the touch using the same rating scale of part 1.

Then, participants were asked to place their left arm palm facing down on a table, with the experimenter sitting in front of them. The experimenter marked two adjacent stroking areas, each measuring 9cm long x 4cm wide, with a washable marker on the hairy skin of their forearm as in previous studies (wrist crease to elbow; Crucianelli et al., 2013, 2016, 2018). Participants were provided with wet wipes at the end of the testing session to remove the marks. The experimenter delivered the touch in the proximal-to-distal direction on these two marked areas using a soft cosmetic make-up brush (Natural hair Blush Brush, N. 7, The Boots Company) at five different speeds: two CT-optimal (3 cm/s and 6 cm/s), one borderline (9 cm/s) and two not CT-optimal (18 cm/s and 27 cm/s). The experimenter was trained to deliver the touch at the exact speed by counting the number of strokes within the tactile stimulation window of 3 seconds (e.g. 1 stroke, 3-seconds-long for the 3 cm/s velocity and 6 strokes, each 0.5-seconds-long for the 18 cm/s velocity). Moreover, stimulation was alternated between the two areas on participant’s forearm to minimize habituation (Crucianelli, Metcalf, Fotopoulou, & Jenkinson, 2013), and because CT fibres are easily fatigued (Vallbo, Olausson, & Wessberg, 1999). Additionally, delivering the touch inside the delineated rectangular spaces allowed the experimenter to be constant not only in space, but also in force/pressure by controlling the lateral spreading of the brush bristles.

Three trials for each speed, for a total of fifteen trials, have been delivered to each participant. The order of conditions was randomized across participants, and after each trial participants used the rating scale ranging from 0 (not at all pleasant) to 100 (extremely pleasant) to judge the pleasantness of the tactile sensation. To avoid any visual feedback of the tactile stimuli or distractions, participants were asked to wear a disposable blindfold throughout the affective touch task.

### 2.4. Design and data analysis

#### Main Analyses

Our task on tactile pleasantness anticipation had a 3 (Group: AN vs. RAN vs. HC) x 2 (Material: soft vs. rough) mixed factorial design, with repeated measures on the latter factor. Participants had to estimate the pleasantness of the material just by imagining it, without having any visual or tactile information about the material or object. The pleasantness score of the two soft and rough materials were then averaged to obtain one pleasantness score for soft and one score for rough touch. Our task on tactile pleasantness perception had a 3 (Group: AN vs. RAN vs. HC) x 3 (Stroking Velocity: Optimal: 3cm/s, 6cm/s vs. Borderline: 9cm/s vs. Suboptimal: 18cm/s, 27cm/s) mixed factorial design, with repeated measures on the latter factor. The pleasantness scores of the two slow (3 and 6cm/s) and the two fast (18 and 27cm/s) velocities were averaged to obtain one value of pleasantness for slow/CT-optimal and one value for fast/CT-non-optimal touch, which were then compared to the borderline velocity and between them. In both the parts of the experiment, the dependent variable was the perceived pleasantness of touch, as measured by means of a pleasantness rating scale ranging from 0 (not at all pleasant) to 100 (extremely pleasant). The scale was presented visually, and participants were responding verbally by choosing a number between 0 and 100.

Statistical analyses were conducted with SPSS version 25.0. The data were tested for normality by means of the Shapiro-Wilk test and found to be non-normal (p < .05). Subsequent Log, Square Root and Reciprocal transformations did not correct for the normality violations, therefore appropriate non-parametric tests were used to analyze the data (described below). Bonferroni-corrected planned contrasts were used to follow up significant effects and interactions. To assess whether there was a difference between groups in the anticipation of pleasantness of different materials, we computed the difference between soft and rough, and run an analysis to check any effect of group on this differential score, and if significant we followed up with Bonferroni-corrected, comparisons on this differential between the HC and the two clinical groups separately. Similarly, to assess whether there was a difference between groups in the perception of affective touch specifically in relation to the CT afferents system, we computed the difference between slow and fast velocities and examine the effect of group on this differential. If significant, we followed up with Bonferroni-corrected comparisons on this differential between HCs and the two clinical groups separately.

#### Exploratory Analyses

In exploratory (due to the relatively small sample size for such analyses) regression analyses, we explored whether anticipatory beliefs about the pleasantness of soft and rough materials and related symtpoms of alexithymia and poor interoceptive sensibility in AN (see below) could predict the perception of CT-optimal and suboptimal touch in each group and their difference. Specifically, we conducted three multiple regressions to explore to what extent the estimated pleasantness of rough and soft materials and EDI, BAQ and TAS scores in each group could predict the perceived pleasantness of CT-optimal touch and separately that of borderline and CT-non-optimal touch.

Next, a multiple regression was run to explore whether the estimated pleasantness of rough and soft materials and EDI, BAQ and TAS scores were significant predictors of tactile sensitivity (difference in pleasantness between slow/CT-optimal touch and fast/CT-non-optimal touch). We conducted this regression in the three groups separately.

## Results

### 2.5. Demographic and mood

Demographic and clinical measures are summarized in Table 2. The three groups did not differ significantly in their age; however, as expected, the BMI was significantly higher in the HC and RAN groups compared to the AN group. In addition, the TAS questionnaire indicated significantly higher alexithymia scores in the AN group compared to HCs and RANs, in line with previous research (e.g. Torres et al., 2015). As expected, the interoceptive awareness’ score of the EDI were significantly higher in people with AN compared to RAN and HC, suggesting an impairment in interoceptive sensibility (Pollatos et al., 2008). A similar pattern was observed in the BAQ questionnaire but there were no significant differences between the groups.

**Table 2.** Summary of demographic and clinical measures. Values provided are means and standard deviation (in parentheses). ** p <.001 difference between groups.

| Group | AGE | BMI | BAQ | EDI | TAS |
| --- | --- | --- | --- | --- | --- |
| AN (27) | 28.40 (10.44) | 14.89 (3.41) | 3.93 (1.01) | 3.68 (1.04) | 60.19 (15.37) |
| RAN (24) | 24.25 (4.51) | 20.15 (2.97) | 4.32 (1.03) | 2.87 (1.02) | 49.29 (15.46) |
| HC (27) | 24.48 (4.61) | 21.39 (3.04) | 4.58 (0.85) | 2.18 (0.59) | 40.04 (7.97) |
| *p* | 0.75 | < 0.01** | 0.05 | < 0.01** | < 0.01** |

### 2.6. Pleasantness ratings

Kolmogorov-Smirnoff test run on the main variables of interest (i.e. all pleasantness scores) showed that both the estimation of pleasantness of materials and the touch pleasantness had a distribution that deviated from normality (all *p*_s_ between 0.001 and 0.06). None of the transformations performed on the variables (log transformation, square root transformation and reciprocal transformation) corrected the data in a way that suited a normal distribution. Therefore, data were analysed using non-parametric tests.

### The Anticipation of Tactile Affectivity

First, the pleasantness scores of the two soft and the two rough materials were averaged to obtain one value for soft and one value for rough materials/objects. To explore the main effect of material, we compared the estimation of pleasantness of soft vs. rough materials, regardless of group. A Wilcoxon Signed Ranks Test revealed a significant main effect of material (Z = −7.35, p < .001), with soft material/objects (M = 67.5; SE = 10.61) been rated as significantly more pleasant than rough materials/objects (M = 10.00; SE = 2.30).

To investigate the main effect of group, we compared the anticipation of pleasantness of the three groups regardless of material. A Kruskal-Wallis Test revealed that group had no effect on anticipation (χ^2^(2) = 3.55; p = 0.169) with a mean rank of anticipation of tactile effectivity of 38.76 for AN group, 33.60 for RAN group and 45.48 for HC group.

Next, to investigate the interaction between group and material, we calculated the differential between the estimation of pleasantness of soft and rough material to obtain one value indicative of the sensitivity of each participant. We then investigated the effect of group on this sensitivity score. A Kruskal-Wallis test showed that group had a significant effect on this difference (χ^2^(2)=15.30, p <.001) with a mean rank of 27.61 for AN group, 39.17 for RAN group and 51.69 for HC group. Post hoc analyses were run by pairwise comparisons with adjusted *p*-values (α= 0.025). Results of Mann-Whitney U tests showed a significant difference in the estimation of pleasantness of soft vs. rough materials between AN patients and HCs (U = 150.50, p < 0.01) but not between RAN and HCs (U = 209, p = 0.029; just a trend).

**Figure 2.**
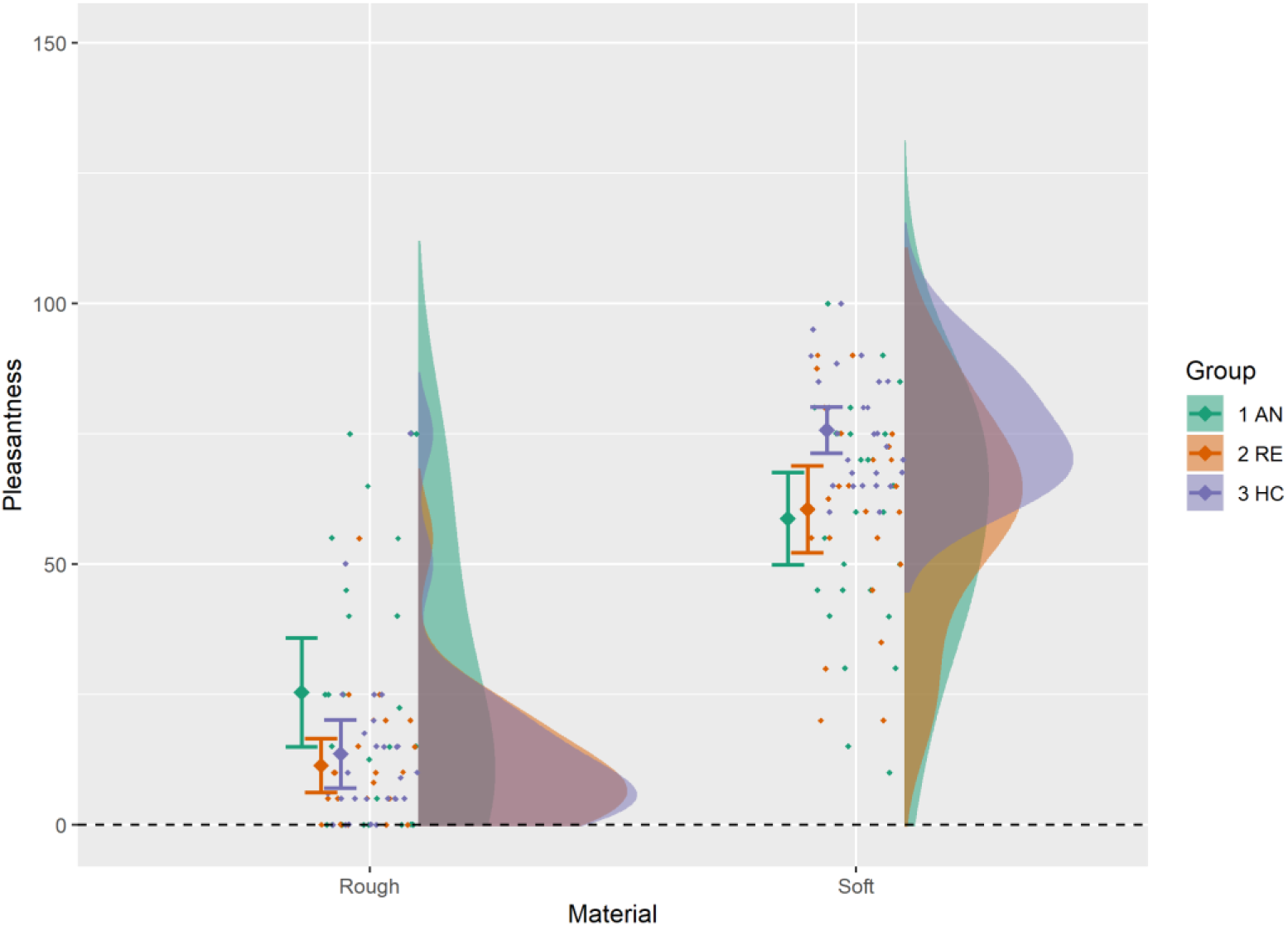
Estimation of pleasantness of soft and rough materials/objects between the three groups: women with anorexia nervosa (AN), women who have recovered from anorexia nervosa (RE) and matched healthy controls (HC).

### The Perception of Tactile Affectivity

To explore the main effect of velocity, we compared the estimation of pleasantness of slow vs. borderline vs. fast velocities, regardless of group. A Friedman Test revealed a significant main effect of velocity (Chi-Square (2) = 78.76, p < .001), with slow (M = 69.0; SE = 2.77) and borderline (M = 73.3; SE = 2.83) touch been rated as significantly more pleasant than fast (M = 44.17; SE = 2.68) touch (see Figure 3), in line with previous studies (Crucianelli et al., 2018). A Kruskal-Wallis test showed a significant main effect of group on the total touch pleasantness ratings (χ^2^(2) = 9.31, p = 0.01), with a mean rank score of 32.93 for AN, 34.85 for RAN and 50.20 for HCs. Bonferroni corrected post-hoc analyses (α = 0.025) revealed that HCs (M = 67.3; SE = 2.90) perceived touch as significantly more pleasant than AN (M = 50.0; SE = 5.03) (Z = −2.60; p = 0.009) and RAN (M = 48.3; SE = 4.50) (Z = −2.61, p = 0.008) regardless of velocity. (see Figure 3). To investigate the interaction between velocity and group, we calculated the differential between the pleasantness scores of slow and fast touch to obtain a value indicative of the CT pleasantness sensitivity of each participant. We then investigated the effect of group on this sensitivity score. A Kruskal-Wallis test showed non-significant interaction between group and velocity (χ^2^(2) = 1.57, p = 0.46), with a mean rank score of 36.69 for AN, 37.75 for RAN and 43.87 for HCs. So, although HCs rated CT-optimal touch as more pleasant than the clinical groups on average, the groups did not differ significantly in how the rated the pleasantness of CT-optimal (slow) vs. non-optimal (fast) touch.

**Figure 3.**
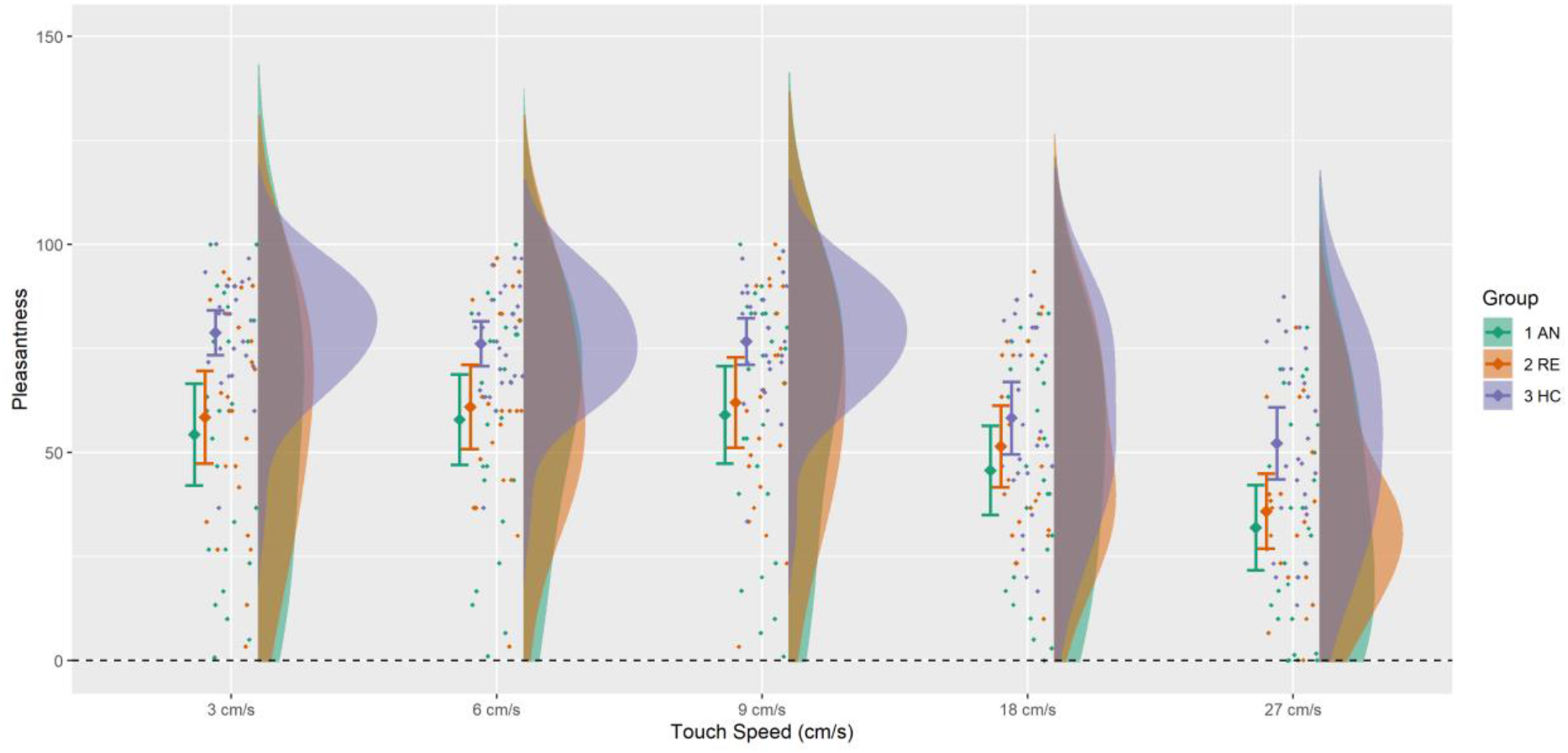
Pleasantness ratings for the five velocities for the three groups: women with anorexia nervosa (AN), women who have recovered from anorexia nervosa (RAN) and matched healthy controls (HC).

We have also run a separate post-hoc analysis on the effect of group on the borderline velocity only, which has been found to be interoceptively relevant (Crucianelli et al., 2018). A Kruskal-Wallis test showed a non-significant main effect of group on the total touch pleasantness ratings (χ^2^(2) = 5.40, p = 0.07), with a mean rank score of 34.67 for AN, 35.75 for RAN and 47.67 for HCs.

### Exploratory analyses

In exploratory (due to the relatively small sample size for such analyses) multiple regression analyses, we explored whether anticipatory beliefs about the pleasantness of soft and rough materials and related symtpoms of alexithymia and poor interoceptive sensibility in AN could predict the perception of CT-optimal and suboptimal touch in each group and their difference. One of the assumptions of regression is that the residuals, rather than the actual variable, must be normally distributed, and this criterion was met.

We ran three multiple regressions to explore whether estimation of pleasantness, TAS, BAQ and EDI were significant predictors of tactile pleasantness at CT-optimal, borderline and CT-non-optimal velocities, separately. We repeated these three regressions for each group. We found a significant regression equation for *CT-optimal touch* in AN (R = 0.746, R^2^ = 0.434, *F*(5, 23) = 4.53, *p* = 0.008) and in RAN (R = 0.766, R^2^= 0.586, *F* (5, 21) =4.53, *p*= 0.009) but not in the HCs group (R = 0.562, R^2^= 0.316, *F* (5, 25) = 1.85, *p*= 0.149). The results for the *borderline velocity*, showed a significant regression equation for RAN (R = 0.718, R^2^ = 0.516, *F*(5, 21) = 3.41, *p* = 0.027). Results for HCs (R = 0.627, R^2^ = 0.394, *F* (5, 25) = 2.60, *p*= 0.058) and AN (R = 0.638, R^2^ = 0.407, *F*(5, 23) = 2.47, *p* = 0.072) showed trends to significance. In terms of *CT-non-optimal touch*, we found a significant regression equation for AN (R = 0.680, R^2^ = 0.462, *F* (5, 23) = 3.09, *p* = 0.035). Results for HCs (R = 0.628, R^2^ = 0.394, *F* (5, 25) = 2.60, *p* = 0.057) and RAN (R = 0.670, R^2^ = 0.449, *F*(5, 21) = 2.61, *p* = 0.066) showed trends to significance. Full results are reported on Table 3.

**Table 3.** Unstandardized coefficient B, standard error of B, standardised coefficient β and p values are reported. * indicates p <0.05 **indicates p < 0.01

| **ANOREXIA NERVOSA** |  |  |  |  |
| --- | --- | --- | --- | --- |
| **CT-optimal touch** | **B** | **SE B** | ***B*** | ***P*** |
| Soft material | 0.34 | 0.24 | 0.26 | 0.18 |
| Rough material | 0.31 | 0.19 | 0.29 | 0.13 |
| EDI | -10.53 | 5.78 | -0.40 | 0.08 |
| BAQ | -11.96 | 4.73 | -0.44 | 0.02* |
| TAS | -0.25 | 0.36 | -0.14 | 0.49 |
| **Borderline** | **B** | **SE B** | ***B*** | ***P*** |
| Soft material | 0.20 | 0.29 | 0.15 | 0.48 |
| Rough material | 0.26 | 0.23 | 0.24 | 0.27 |
| EDI | -6.07 | 6.82 | -0.22 | 0.39 |
| BAQ | -11.46 | 5.58 | -0.42 | 0.27 |
| TAS | -0.48 | 0.42 | -0.27 | 0.27 |
| **CT-non-optimal touch** | **B** | **SE B** | ***B*** | ***P*** |
| Soft material | 0.01 | 0.24 | 0.01 | 0.95 |
| Rough material | 0.34 | 0.19 | 0.35 | 0.10 |
| EDI | -6.73 | 5.73 | -0.28 | 0.26 |
| BAQ | -6.42 | 4.69 | -0.27 | 0.19 |
| TAS | -0.54 | 0.35 | -0.34 | 0.14 |
| **Tactile sensitivity** | **B** | **SE B** | ***B*** | ***P*** |
| Soft material | 0.32 | 0.21 | 0.38 | 0.13 |
| Rough material | -0.03 | 0.16 | -0.04 | 0.87 |
| EDI | -3.80 | 4.91 | -0.22 | 0.45 |
| BAQ | -5.54 | 4.02 | -0.32 | 0.18 |
| TAS | 0.29 | 0.30 | 0.26 | 0.34 |
| **RECOVERED ANOREXIA NERVOSA** |  |  |  |  |
| **CT-optimal touch** | **B** | **SE B** | ***B*** | ***P*** |
| Soft material | 0.07 | 0.03 | 0.47 | 0.02* |
| Rough material | 0.19 | 0.40 | 0.10 | 0.64 |
| EDI | 15.64 | 9.30 | 0.66 | 0.11 |
| BAQ | 15.40 | 4.50 | 0.65 | 0.003** |
| TAS | -0.47 | 0.61 | -0.30 | 0.45 |
| **Borderline touch** | **B** | **SE B** | ***B*** | ***P*** |
| Soft material | 0.07 | 0.03 | 0.44 | 0.04* |
| Rough material | 0.20 | 0.47 | 0.10 | 0.66 |
| EDI | 10.94 | 10.70 | 0.44 | 0.32 |
| BAQ | 15.60 | 5.17 | 0.61 | 0.008** |
| TAS | -1.23 | 0.63 | -0.75 | 0.07 |
| **CT-non-optimal touch** | **B** | **SE B** | ***B*** | ***P*** |
| Soft material | 0.04 | 0.03 | 0.33 | 0.14 |
| Rough material | -0.32 | 0.42 | -0.18 | 0.45 |
| EDI | 23.11 | 9.66 | 1.09 | 0.03* |
| BAQ | 12.48 | 4.67 | 0.58 | 0.02* |
| TAS | -1.42 | 0.56 | -1.01 | 0.02* |
| **Tactile sensitivity** | **B** | **SE B** | ***B*** | ***P*** |
| Soft material | 0.02 | 0.02 | 0.30 | 0.21 |
| Rough material | 0.51 | 0.30 | 0.44 | 0.11 |
| EDI | -7.46 | 6.96 | -0.54 | 0.30 |
| BAQ | 2.92 | 3.36 | 0.21 | 0.40 |
| TAS | -0.07 | 0.41 | -0.08 | 0.86 |
| **HEALTHY CONTROLS** |  |  |  |  |
| **CT-optimal touch** | **B** | **SE B** | ***B*** | ***P*** |
| Soft material | 0.45 | 0.27 | 0.38 | 0.10 |
| Rough material | 0.06 | 0.15 | 0.07 | 0.70 |
| EDI | 6.11 | 5.20 | 0.27 | 0.25 |
| BAQ | -1.84 | 3.70 | -0.12 | 0.62 |
| TAS | -0.63 | 0.42 | -0.38 | 0.15 |
| **Borderline** | **B** | **SE B** | ***B*** | ***P*** |
| Soft material | 0.58 | 0.27 | 0.45 | 0.04* |
| Rough material | 0.08 | 0.15 | 0.09 | 0.62 |
| EDI | 7.29 | 5.63 | 0.30 | 0.21 |
| BAQ | -3.09 | 3.79 | -0.18 | 0.42 |
| TAS | -0.68 | 0.43 | -0.37 | 0.13 |
| **CT-non-optimal touch** | **B** | **SE B** | ***B*** | ***P*** |
| Soft material | 0.72 | 0.41 | 0.37 | 0.09 |
| Rough material | 0.11 | 0.23 | 0.08 | 0.65 |
| EDI | 15.83 | 8.46 | 0.43 | 0.08 |
| BAQ | -4.87 | 5.70 | -0.19 | 0.40 |
| TAS | -0.04 | 0.65 | -0.02 | 0.95 |
| **Tactile sensitivity** | **B** | **SE B** | ***B*** | ***P*** |
| Soft material | -0.27 | 0.32 | -0.18 | 0.41 |
| Rough material | -0.05 | 0.18 | -0.05 | 0.79 |
| EDI | -10.44 | 6.70 | -0.35 | 0.13 |
| BAQ | 3.02 | 4.15 | 0.15 | 0.51 |
| TAS | -0.68 | 0.49 | -0.05 | 0.18 |

Next, we run three multiple regressions with beliefs of pleasantness, TAS, BAQ and EDI as predictors and the differential between slow and fast (tactile sensitivity) as outcome measure in the AN, RAN and HCs group separately. In the AN group, results showed a non-significant regression equation (R = 0.482, R^2^ = 0.233, *F*(5, 23) = 1.09, *p*= 0.398). Similarly, we found a non-significant regression equation for the RAN group (R = 0.579, R^2^ = 0.335, *F*(5, 21) = 1.61, *p* = 0.213). However, we found a trend to significance for the equation in the HCs group (R = 0.629, R^2^ = 0.396, *F*(5, 25) = 2.62, *p* = 0.056, see Table 3).

## 3. Discussion

This study aimed to investigate (1) whether the anticipation and perception of pleasant touch would be reduced in patients who have recovered from AN as well as in patients with acute AN compared to healthy controls; (2) whether this tactile anhedonia will be best explained by deficits in the CT-afferent system, top-down beliefs about the pleasantness of affective touch, or their combination. The results confirmed our first prediction, showing that individuals with and recovered from AN anticipated and perceived affective touch as overall less pleasant compared to healthy controls. However, contrary to our predictions and two previous studies (Crucianelli et al., 2016; Davidovic et al., 2018), the reduction in the perceived tactile pleasantness was not specific to the CT-optimal (i.e. slow) stroking velocities but was also observable to a lesser degree in non-CT optimal touch. Also, contrary to our prediction there was a group by material interaction in the anticipation of affective touch, mainly driven by the fact that the HCs showed a larger than the AN group differentiation between how pleasant they expect soft materials to feel in comparison to rough materials, with the difference between HCs and RAN not reaching significant levels. Interesting, exploratory analyses revealed that these anticipatory beliefs predicted individual differences in the perceived pleasantness of CT-optimal but not non-CT optimal touch within these two latter groups (in addition to alexithymia and interoceptive sensibility in the RAN group), but not in the AN group. Instead, individual differences in the perceived pleasantness of CT-optimal in the AN group were predicted only by interoceptive sensibility scores. We discuss these findings in detail below.

Our results regarding the reduced perceived pleasantness of gentle touch in individuals with and recovered from AN are consistent with long-standing findings regarding anhedonia in AN, now extending them to the tactile domain. People with AN show reduced pleasure for a range of stimuli, such as food (Davis & Woodside, 2002), amusing stimuli and sexual activity (Tchanturia, et al., 2012). Anhedonia also correlates with strenuous physical exercise in this population (Davis & Woodside, 2002) and several studies have suggested that the reduced reward sensitivity in AN is coupled with an increase tendency for cognitive and behavioural control, including the control of one’s energy intake (Wagner et al., 2007; Kaye al., 2013). Moreover, the fact that this tactile anhedonia was observed in both recovered, as well as acute patients with AN suggests that the symptom is not a secondary consequence of starvation, but it may be a trait that persists even after recovery.

However, contrary to our prediction, we did not find evidence to support the idea that the observed tactile anhedonia was related to abnormalities of the CT system, a tactile modality thought to be mediated peripherally by unmyelinated, slow-conducting C-tactile (CT) afferents (Löken et al., 2009) and to be centrally processed by an interoceptive pathway converging at the insular cortex (Morrison, 2016; Kirsch et al., 2020). Although our clinical groups showed reduced perceived pleasantness for CT-optimal stimulation in comparison to healthy volunteers, this finding applied for non-CT-optimal touch too. More broadly, while microneurography and psychophysical investigations have indicated that C-tactile afferents have peak firing rates that positively correlate with the perceived touch pleasantness of gentle touch (Löken et al., 2009), a recent study showed that spinothalamic ablation in humans had no effect on pleasant touch perception, suggesting a more integrated hedonic and discriminative spinal processing from the body (Marshall et al., 2019). Thus, future studies in AN need to explore further tactile and interoceptive modalities in order to draw firm conclusions about the mechanisms that cause tactile anhedonia in this population.

A further unexpected finding in our study may begin to address such complexity in how the affectivity of touch is perceived in AN. Specifically, patients with acute AN, and in non-significant levels also individuals recovered from AN, showed reduced pleasantness differentiation in how they imagined soft versus rough material touching their skin. In other terms, the sensory anhedonia observed in this study may not be a mere matter of how bottom-up sensory signals travel to the brain, but may also be affected by abnormalities in top-down, expectations. This is consistent with a recent study finding both behavioural and neural abnormalities in how a small sample of RAN patients anticipated CT-optimal touch (Bischoff-Grethe et al., 2018). More broadly, this finding is consistent with the idea that in AN (Khalsa et al., 2015) there is a dysregulated ability to adequately predict and cognitively represent and regulate what is happening in the physiological state of the body, resulting in symptoms such as alexithymia (Shah et al., 2016).

Indeed, our exploratory analyses revealed that while the perceived pleasantness of CT-optimal touch in HCs was predicted by their ability to differentiate the sensory consequences of different tactile materials, the ratings of patients with acute AN were predicted only by their interoceptive sensibility scores, while those of the RAN group were predicted by both factors. These results further suggest that tactile anhedonia in AN may be secondary to impairments of bodily signals identification, expression and regulation, but given the relatively small size of our samples these findings will need to be explored further in future studies.

In conclusion, this study found that the anticipation and perception of pleasant touch is reduced in patients who have recovered from AN as well as in patients with acute AN compared to healthy controls, but this tactile anhedonia is not best explained by deficits in the CT-afferent system. It appears that top-down beliefs about the pleasantness of affective touch are also affected in this disorder and tactile anhedonia is best explained in reference to concomitant symptoms of interoceptive sensibility.

## Authors Contributions

LC, BD and AF conceived and designed the experiment. LC, BD, AS, DG and VN performed data collection. OG, SB and PT supported the clinical recruitment of individuals with and recovered from Anorexia Nervosa. LC analysed the data, under supervision of AF. LC, BD, AS and AF wrote the manuscript. All the authors approved the manuscript before submission.

## Acknowledgements

We thank all the patients for their kindness and willingness to take part in the study. Authors would like to thank Ilia Galouzidi for assistance with data entering. The study was supported by a “European Research Council Consolidator Award” [ERC-2018-COG-818070] to A.K. and a Neuropsychoanalysis Foundation grant to L.C.

